# Protein kinase Cε activation induces EHD-dependent degradation and downregulation of K_ATP_ channels: Implications for glucose stimulated insulin secretion

**DOI:** 10.1101/2020.01.15.907386

**Authors:** Christopher J Cockcroft, Paul Manna, Rucha Karnik, Tarvinder K Taneja, David Wrighton, Jamel Mankouri, Hong-Lin Rong, Asipu Sivaprasadarao

**Affiliations:** School of Biomedical Sciences, Faculty of Biological Sciences, University of Leeds, LS2 9JT, Leeds, U.K; Multidisciplinary Cardiovascular Research Centre, University of Leeds, LS2 9JT, Leeds, U.K

**Keywords:** K_ATP_ channels, Protein kinase Cε, EHD, Type 2 diabetes, β-cells, dileucine motif

## Abstract

Pancreatic β-cells have the unique ability to couple glucose metabolism to insulin secretion. This capacity is generally attributed to the ability of ATP to inhibit K_ATP_ channels, and the consequent β-cell membrane depolarization and excitation. This notion has recently been challenged by a study which demonstrated that high glucose (HG) downregulates the cell surface K_ATP_ channels, and thereby leads to β-cell depolarisation and excitation. The authors attributed the downregulation to HG-induced protein kinase C (PKC) activation and the consequent increase in channel endocytosis. This interpretation, however, is inconsistent with our previous findings that PKC activation does not affect endocytosis. To address this controversy, we revisited the problem: we have used cell biological and electrophysiological approaches combined with the pharmacological activator of PKC, PMA (phorbol 12-myristate 13-acetate). We first confirm that PKC does not play a role in K_ATP_ channel endocytosis; instead, it downregulates the channel by promoting lysosomal degradation coupled with reduced recycling. We then show that (i) mutation of the dileucine motif (^355^LL^356^) in the C-terminal domain of the Kir6.2 subunit of the K_ATP_ channel complex prevents lysosomal degradation; (ii) lysosomal targeting is mediated by the EHD (Eps15 homology domain– containing) proteins; and (iii) the PKC isoform responsible for channel degradation is PKCε. Taken together with the published data, we suggest that HG promotes β-cell excitability via two mechanisms: ATP-dependent channel inhibition and ATP-independent, PKCε-dependent channel degradation. The results likely have implications for glucose induced biphasic insulin secretion.

## Introduction

Pancreatic β-cells have the remarkable ability to couple glucose metabolism to insulin secretion. This ability is, at least in part, conferred by the plasma membrane ATP-sensitive potassium (K_ATP_) channels (Ashcroft and Rorsman, 2013; Nichols, 2006; Seino and Miki, 2003). When blood glucose levels rise, glucose uptake and metabolism by the pancreatic β-cell increase, resulting in an increase in the cytosolic [ATP]/[ADP] ratio. Rise in the [ATP]/[ADP] ratio leads to a cascade of events including closure of K_ATP_ channels, membrane depolarisation (due to reduced K^+^ efflux), activation of voltage-gated calcium channels, and Ca^2+^ influx. The rise in cytosolic Ca^2+^ ultimately triggers insulin release (Ashcroft, 2006; Ashcroft and Rorsman, 2013; Nichols, 2006).

Structure-function relationship studies (Ashcroft, 2006; Ashcroft and Rorsman, 2013; Nichols, 2006)and the more recent structural data (Lee et al., 2017; Li et al., 2017; Martin et al., 2017) provide a mechanistic insight into how the [ATP]/[ADP] ratio controls K_ATP_ channel gating. K_ATP_ channels are made up of two types of subunit: the inwardly rectifying K^+^ channel Kir6.2 (encoded by *KCNJ11*) and ABC transporter SUR1 (encoded by *ABCC8*) (Inagaki et al., 1995; Nichols, 2006). Kir6.2 subunits are assembled into a tetramer forming a pore for K^+^ permeation as well as binding sites for the inhibitory ATP. Surrounding the Kir6.2 tetramer are four SUR1 subunits that harbour binding sites for the activating nucleotide, Mg-ADP. The reciprocal functional effects of ATP and ADP endows the K_ATP_ channel with the ability to couple glucose metabolism to β-cell excitability and insulin secretion (Ashcroft and Rorsman, 2013; Nichols, 2006).

Researchers have long believed that ATP inhibition of K_ATP_ channels represents the dominant mechanism linking glucose metabolism to insulin secretion (Ashcroft and Rorsman, 2013; Nichols, 2006; Seino and Miki, 2003). However, a recent study challenged this notion (Han et al., 2018). The authors proposed that downregulation of K_ATP_ channels plays a greater role than ATP-dependent gating in β-cell excitability. In support of this, they demonstrated that high glucose (HG) causes profound downregulation of plasma membrane K_ATP_ channels, leading to membrane depolarisation and β-cell excitation. These HG effects were prevented by dynasore, an inhibitor of endocytosis, as well as inhibition of protein kinase C (PKC). This has led the authors to conclude that PKC-induced endocytosis of K_ATP_ channels drives glucose-stimulated β-cell excitation (Han et al., 2018). While the ability of PKC to downregulate K_ATP_ channels is consistent with a number of other studies (Bruederle et al., 2011; Hu et al., 2003), including ours (Mankouri et al., 2006; Manna et al., 2010), the mechanism proposed by the authors is inconsistent with our previous studies (Manna et al., 2010). We have shown that K_ATP_ channels undergo constitutive endocytosis and that PKC does not affect the rate of endocytosis. Our data suggested that downregulation is caused by the increased degradation of the endocytosed channels (Manna et al., 2010). To address this controversy, we revisited the problem using genetic tools in conjunction with pharmacological reagents. First, we introduced mutations into Kir6.2 to inactivate the dileucine trafficking motif, ^355^LL^356^, because this motif is essential for PKC-mediated downregulation of the channel (Hu et al., 2003; Mankouri et al., 2006). Second, we used a mutant of EHD-1, a protein with an established role in recycling of many membrane proteins (Naslavsky and Caplan, 2011; Park et al., 2004; Picciano et al., 2003). Third, we have used dominant negative constructs of PKC isoforms to complement the pharmacology and to underpin the isoform responsible channel downregulation. Our results provide new mechanistic insights into how PKC downregulates the plasma membrane K_ATP_ channels. We discuss the possible relevance of our findings to glucose stimulated biphasic insulin secretion (GSIS).

## Results

### Mutation of the dileucine motif in Kir6.2 abrogates PMA-induced inhibition of K_ATP_ channel recycling

Although the ability of PKC to downregulate plasma membrane K_ATP_ channels is fairly well established, there is controversy surrounding the underlying mechanism. Hu et al reported that PKC stimulates K_ATP_ channel endocytosis, and that the effect is mediated by the dileucine traffic motif on Kir6.2 (^355^LL^356^) (Hu et al., 2003). However, later studies demonstrated that K_ATP_ channel endocytosis is PKC independent, but confirmed that the dileucine motif is essential for channel downregulation (Mankouri et al., 2006; Manna et al., 2010). Here we asked whether the dileucine motif plays a role in PKC dependent K_ATP_ channel downregulation by affecting their rates of recycling and/ or degradation.

For this, we introduced the LL/AA double mutation into HA (hemaglutinin A)-tagged Kir6.2 (HA-Kir6.2), because the extracellular location of the HA-epitope enables investigation of channel trafficking (Hu et al., 2003; Mankouri et al., 2006; Manna et al., 2010). We have also made a second mutant by substituting a proline for L355 (L355P). This mutation is expected to disrupt the α-helix in which the dileucine motif resides and have an effect similar to that of LL/AA. We have used PMA (phorbol 12-myristate 13-acetate) to investigate the effect of PKC activation on endosomal trafficking and degradation.

We co-expressed the wild-type and mutant HA-Kir6.2 subunits with SUR1 in HEK-293 cells. For simplicity, hereafter, we refer to the HA-Kir6.2-SUR1 as WT (wild-type), the mutants as LL/AA and L355P. As expected, the L355P mutation, like the LL/AA mutation (Hu et al., 2003; Mankouri et al., 2006), increased the steady-state cell surface density of the channel (70.7 ± 9.3% for L355P vs 39.4 ± 8.2 % for WT; *p* < 0.05; Figure 1A-B). As reported previously with the LL/AA mutation (Mankouri et al., 2006); also see Figure 1G), the L355P mutation did not significantly alter the rate of endocytosis (48.9 ± 6.0 % for WT vs 57. 5 ± 13.2 % for L355P; *p* > 0.05) (Figure 1C-D). We examined recycling in the absence and presence PMA. The results show that PMA inhibits recycling of WT, but not the mutant channel (44.4 ± 8 % for WT, *p* < 0.05; 95.8 ± 25 % for L355P; *p* > 0.05) (Figure 1E-F). Similar effects on recycling were also observed with the LL/AA mutant (Figure 1H). These results confirm that PMA-induced downregulation of K_ATP_ channels is not due to increased endocytosis, but due to abrogation of the ability of PMA to inhibit recycling of internalised channels. Importantly, they demonstrate that the inhibitory effect of PMA on channel recycling is mediated by the dileucine motif on Kir6.2.

**Figure 1.**
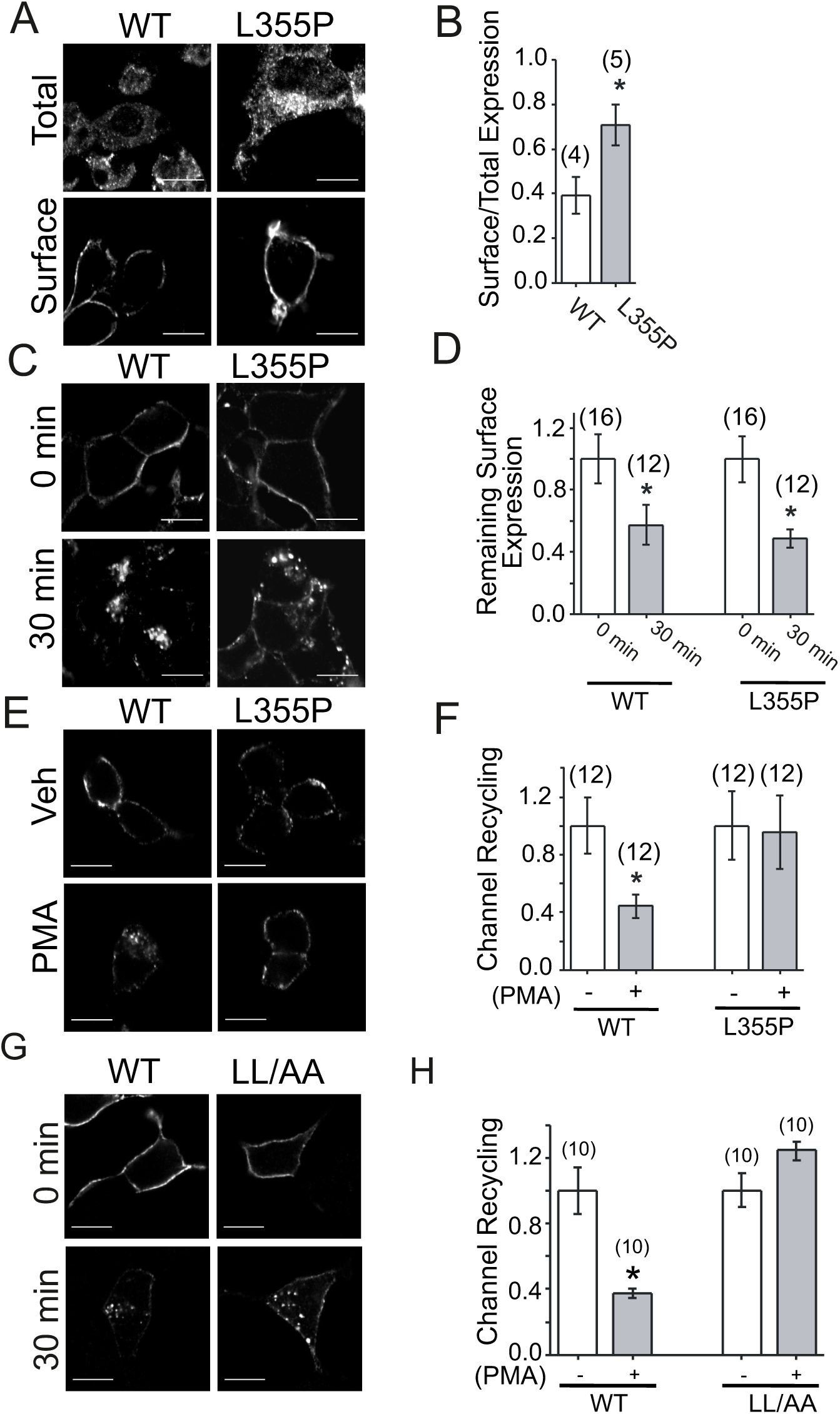
Mutation of the dileucine motif in Kir6.2 increases the K_ATP_ channel surface expression by preventing PKC-mediated inhibition of recycling. HEK-293 cells transfected with the HA-tagged WT or L355P mutant channels were used for panels A-F. (A-B), L355P mutation in Kir6.2 increases the surface expression of recombinant K_ATP_ channels relative to total channels. (A) Representative images of immunostained channels in permeabilised (total) and unpermeabilised (surface) cells. (B) Bar chart showing mean ± SEM of surface density of the L355P mutant channel, expressed as a fraction of total channel protein. (C-D) The L355P mutation does not affect the internalisation rate of the channel. Representative images of WT and L355P mutant channels before (0 min) and after 30 min incubation with anti-HA antibodies at 37 °C (C) and the corresponding mean ± SEM data (D); data were normalised to ‘0’ time. (E-F) PMA reduced recycling of WT, but not the L355P mutant channel. Representative images of internalised WT and L355P mutant channels that have recycled back to the cell surface during a 30 min incubation at 37 °C in the absence (0.1 % DMSO, Veh) or presence of PMA (100 nM/0.1% DMSO) (E), and the corresponding mean ± SEM data (F); data were normalised to 0.1 % DMSO (-) values. (G) LL/AA mutation does not inhibit internalisation. Representative images of HEK-293 cells transfected with the HA-tagged WT or LL/AA mutant channels treated as for C. (H) LL/AA mutation mitigates the ability of PMA to inhibit recycling. Mean ± SEM of recycling of WT or LL/AA mutant channels determined as for F. Numbers at the top of each bar indicate ‘n’; * indicates significance (*p* < 0.05).

### L355P mutation prevents PMA-induced degradation of the endocytosed K_ATP_ channel

The L355P mutation is a missense mutation found in an Afro-Caribbean patient with type 2 diabetes (Sakura et al., 1996). The mutation did not affect the sensitivity of the channel to metabolic inhibition, leading the authors to conclude that this mutation is unlikely to be a major cause of the disease (Sakura et al., 1996). However, our finding that the L355P mutation upregulates the channel (Figure 1) raises the possibility that altered trafficking could play a causative role in the disease. Thus we focussed on this mutation for the rest of our studies. Dileucine motifs are known play a wide variety of roles in endosomal trafficking including sorting of endocytosed membrane proteins to lysosomes (Bonifacino and Traub, 2003). We have previously shown that PMA treatment increases lysosomal trafficking and degradation of K_ATP_ channels (Manna et al., 2010). We therefore asked whether the L355P mutation abrogates PMA’s ability to promote lysosomal trafficking. Our results indeed show that PMA treatment significantly (*p*<0.05) increased the localisation of the endocytosed WT channels, but not the L355P mutant channels, to CD63-positive lysosomal/late endosomal compartments (Figure 2A-B). We next tested the effect of the L355P mutation on PMA-induced channel degradation using a biotinylation-pulldown assay (Manna et al., 2010). We labelled the cell surface proteins with a membrane impermeable biotinylation reagent, treated the cells with vehicle or PMA for 2 hours (to allow channel degradation), pulled down the HA-tagged (WT and the mutant) channels with Neutravidin-agarose beads and estimated the amount of undegraded HA-Kir6.2 in the pelleted beads by western blotting. Figure 2C shows that PMA treatment caused a significant decrease in the amount of WT (*p*<0.05), but not the L355P mutant subunit (*p*>0.05). These data suggest that the L355P mutation inhibits the ability of PMA to induce K_ATP_ channel degradation. Commensurate with this, PMA treatment significantly decreased the surface levels of the WT (*p*<0.05), but not the mutant channels (*p*>0.05) (Figure 2D). Together, these data indicate that the L355P mutation abolishes the ability of PMA to downregulate the plasma membrane K_ATP_ channels by preventing their lysosomal trafficking and degradation.

**Figure 2.**
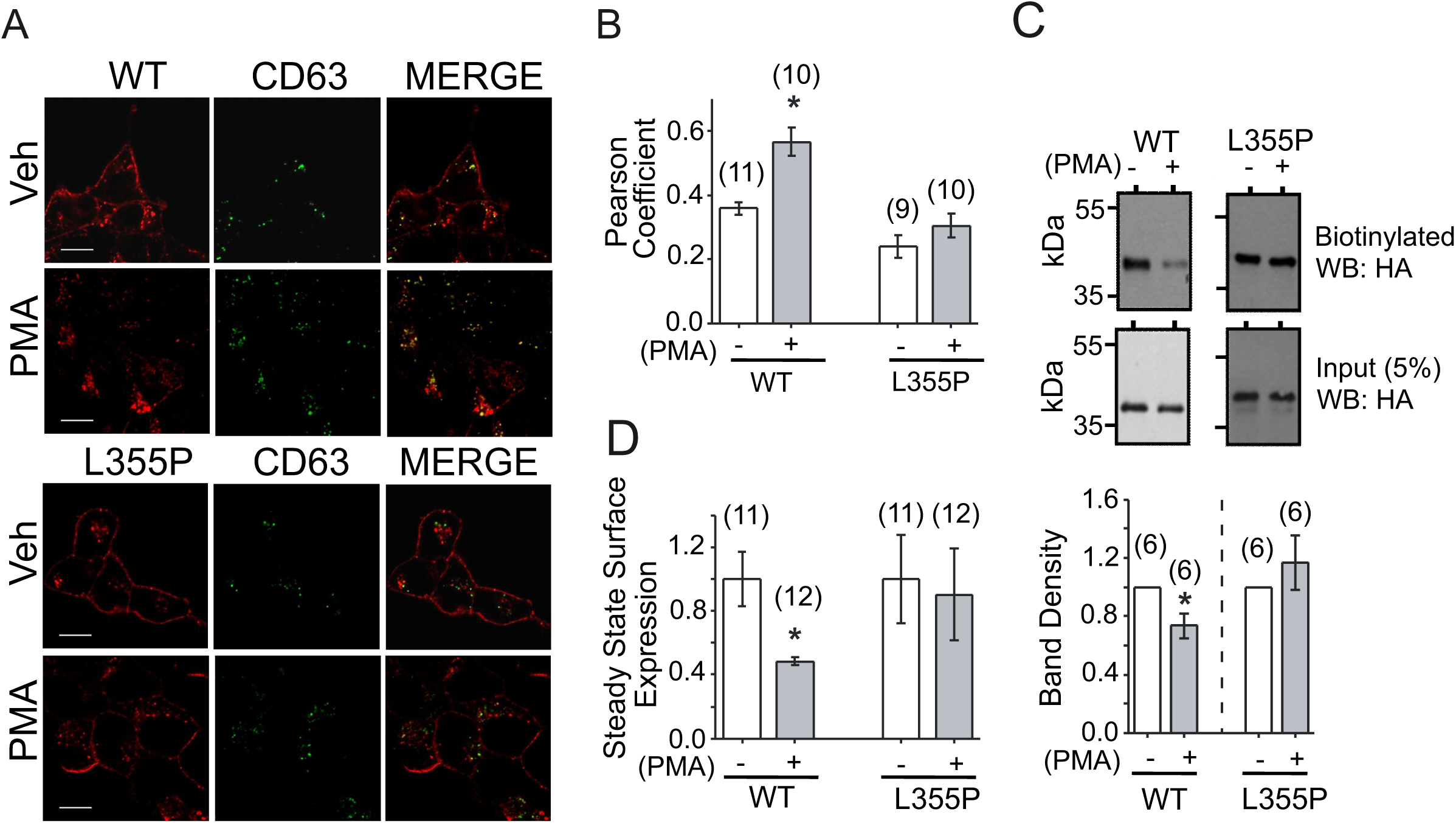
The L355P mutation in Kir6.2 prevents PKC-induced channel degradation. (A-B) PMA increases lysosomal trafficking of internalised WT, but not L355P mutant channels. (A) Representative images (individual and merged) of transfected HEK293 cells stained for internalised K_ATP_ channels and CD63, a marker for lysosomes/late endosomes. Cells were allowed to take up anti-HA antibodies at 37 °C for 30 min in the absence (0.1 % DMSO; Veh) or presence of PMA (100 nM/0.1% DMSO), and labelled with rat anti-HA (Cy3 secondary) and mouse anti-CD63 (Alexa488 secondary) antibodies. (B) Bar chart showing quantification of co-localisation between the internalised channels and CD63; the degree of co-localisation is expressed in terms of Pearson co-efficient. (C) PMA increases the degradation of WT, but not the L355P mutant channels. Following surface biotinylation, cells were treated with either 0.1 % DMSO (-) or 100 nM PMA (+) for 2 hrs at 37 °C. Biotinylated channels were pulled down with NeutrAvidin beads, and detected by western blotting using anti-HA antibodies. Representative blots are shown. The bar chart (bottom panel) shows mean ± SEM of band intensities; data were normalised to respective DMSO (-) treated cells. (D) PMA treatment causes a reduction in the steady-state surface levels of WT, but not the L355P mutant channels; data were normalised to the respective DMSO (-) treated cells. Numbers at the top of each bar in all bar charts indicates ‘n’; * indicates significance (*p* < 0.05).

### EHD-1 proteins promote PKC-mediated degradation of the wild-type, but not L355P mutant channels

There can be two possible explanations for the PMA effect on recycling and degradation. PMA could inhibit recycling and thereby increase the endosomal pool of channels available for lysosomal targeting. Alternatively, PMA could increase lysosomal trafficking and thereby reduce the pool for recycling. EHD-1 (Eps15 homology domain -containing protein-1) is known to play a major role in recycling of membrane proteins (Naslavsky and Caplan, 2011; Park et al., 2004; Picciano et al., 2003). Thus, we first asked whether the internalised K_ATP_ channels traffic to EHD-positive recycling compartments. Immunostaining with pan anti-EHD antibodies showed co-localisation of internalised HA-Kir6.2-SUR1 channels with native EHD proteins (Figure 3A). The extent of co-localisation was greater in PMA-treated cells than the untreated cells and cells co-treated with the PKC inhibitor, chelerythrine (Figure 3A). These data are consistent with a previous report that internalised K_ATP_ channels traffic to the EHD-positive compartments (Hu et al., 2003), suggesting a potential role for EHD proteins in the channel recycling.

**Figure 3.**
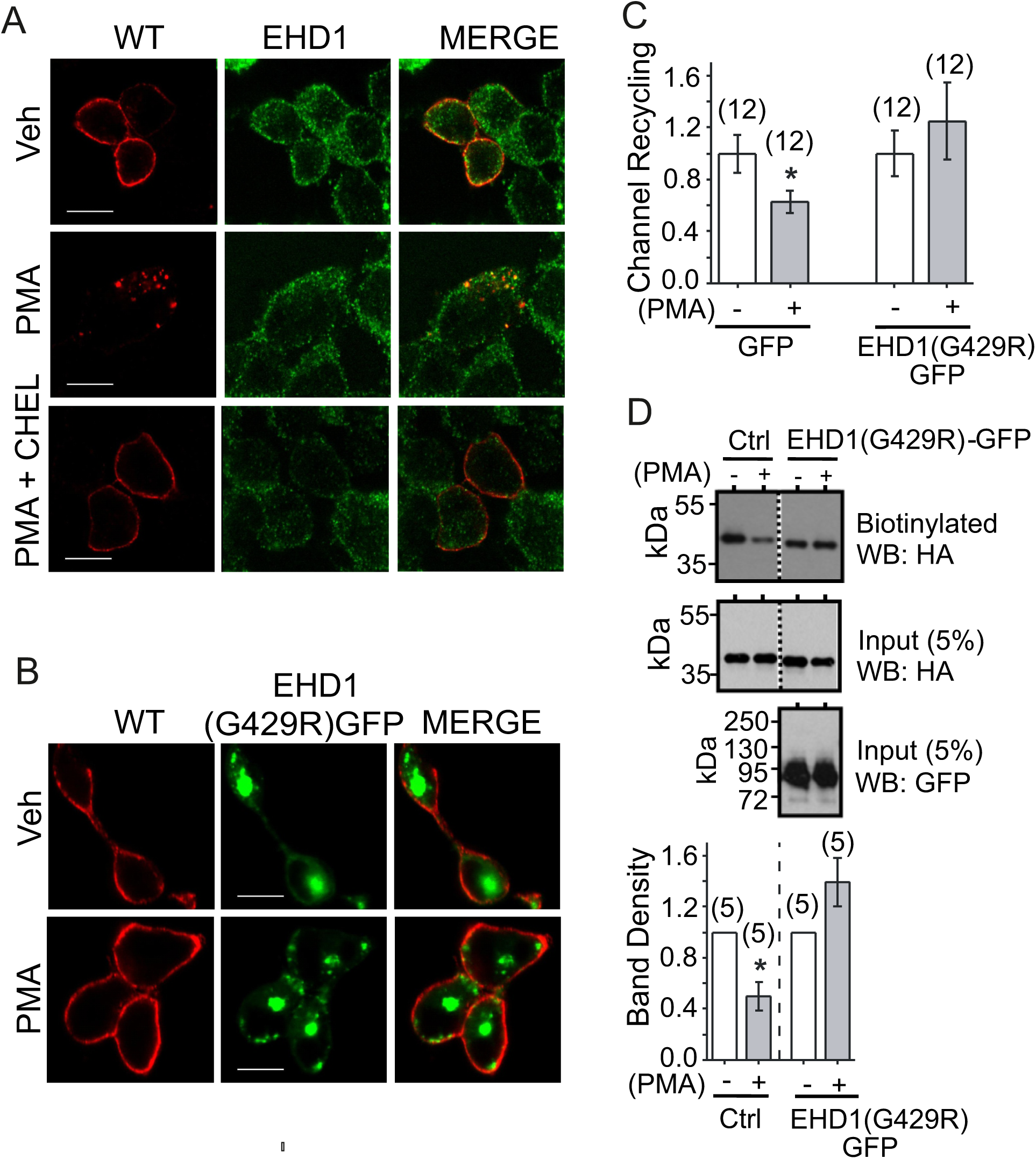
Dominant negative EHD-1 prevents PKC-induced inhibition of recycling and degradation of wild-type K_ATP_ channels. (A) Internalised K_ATP_ channels co-localise with endogenous EHD-1. HEK 293 cells expressing the WT channels were incubated at 37 °C for 30 min with anti-HA antibodies in the absence (Veh, 0.1 % DMSO), or presence of PMA (100 nM), or PMA (100 nM) plus chelerythrine (10 µM) before staining for the channel and endogenous EHD-1. Representative individual and merged images are shown. (B) Dominant negative EHD-1(G249R)-GFP prevents PMA induced endosomal accumulation of WT K_ATP_ channels. HEK 293 cells co-expressing the channel and EHD-1(G249R)-GFP were allowed to internalise anti-HA antibodies in the absence (Veh, 0.1 % DMSO), or presence of PMA (100 nM) and stained with Cy3-conjugated secondary antibodies after permeabilisation. Representative images (individual and merged) are shown. (C) EHD-1(G249R)-GFP rescues PMA-induced reduction in recycling. Cells expressing the WT channels were transfected with either GFP or EHD-1(G429R)-GFP and the effect of PMA on the rate of recycling was determined by as described in the legend for Fig 1F. (D) EHD-1(G249R)-GFP prevents PMA-induced degradation of surface channels. Cells expressing the WT channels were transfected with either GFP (control) or EHD-1(G429R)-GFP and surface biotinylated; degradation of biotinylated Kir6.2-HA was determined as described in the legend to Figure 2C. Representative blots (top panels) and mean ± SEM data (bottom panel) are shown. Numbers at the top of each bar in all bar charts indicates ‘n’; * indicates significance (*p* < 0.05).

To study recycling of membrane proteins, researchers have often used a dominant negative construct of EHD-1, EHD-1(G429R)-GFP (Naslavsky and Caplan, 2011; Park et al., 2004; Picciano et al., 2003). Its ability to inhibit recycling can be readily revealed by an increase in the co-localisation of internalised proteins with EHD-1(G429R)-GFP and a concomitant decrease in the surface levels. We thus expressed EHD-1(G429R)-GFP in cells stably expressing HA-Kir6.2-SUR1. Contrary to the expectation, we found little co-localisation of internalised K_ATP_ channels with EHD-1(G429R)-GFP (Figure 3B). Instead, the majority of channel staining was apparent at the plasma membrane (Figure 3B). These results were surprising given the fact that EHD-1 is known to play a major role in endosomal recycling. To confirm this unexpected result, we examined the effect of EHD-1(G429R)-GFP on recycling. The results show that in GFP-transfected control cells, as expected, PMA significantly inhibited the channel recycling (*p*<0.05); by contrast, in EHD-1(G429R)-GFP transfected cells, PMA failed to inhibit recycling (*p*>0.05) (Figure 3C). Thus our results show that EHD-1(G429R)-GFP is capable of rescuing the apparent inhibitory effect of PMA on K_ATP_ channel recycling.

Studies have shown that EHD-1 is a member of the EHD family comprising four members (1-4) and that EHD-1 can associate with other members the family to form heteromers capable of influencing other endocytic routes, including late endosomal targeting (Naslavsky and Caplan, 2011). Thus we examined the effect of EHD-1(G429R)-GFP on K_ATP_ channel degradation using the biotinylation-based assay. The results indeed demonstrated robust rescue of PMA-induced channel degradation by EHD-1(G429R)-GFP (Figure 3D). Taken together, our results indicate that, by promoting PMA-induced degradation of the channel, EHD proteins downregulate the number of K_ATP_ channels at the plasma membrane.

### EHD-1(G429R)-GFP prevents PMA-induced downregulation of K_ATP_ channel currents in the INS-1 832/13 model pancreatic β-cell line

We next tested the role of EHD in the more physiologically relevant rat INS-1 pancreatic β-cell line. This cell line responds to HG and secretes insulin. Furthermore, HG stimulates PKC activity leading to K_ATP_ channel downregulation as well as depolarisation and electrical excitation of INS-1 cells (Han et al., 2018). Thus we examined the effect of expression of EHD-1(G428R)-GFP on PKC regulation of K_ATP_ currents in this cell line. Whole cell patch clamp recordings show that PMA treatment caused a marked downregulation of K_ATP_ current density (2.56 ± 0.44 nS/pF for control vs 0.73 ± 0.10 nS/pF for PMA; *P* =0.002) in GFP-expressing (control) cells. By contrast, in cells expressing EHD-1(G429R)-GFP, PMA showed no significant effect (*P* =0.04) on the current density (3.23 ±0.63 nS/pF for control vs 2.5 ± 0.68 nS/pF for PMA) (Figure 4A-C). These results indicate that EHD proteins play a role in PMA-induced downregulation of native plasma membrane K_ATP_ channels in the pancreatic β-cell line.

**Figure 4.**
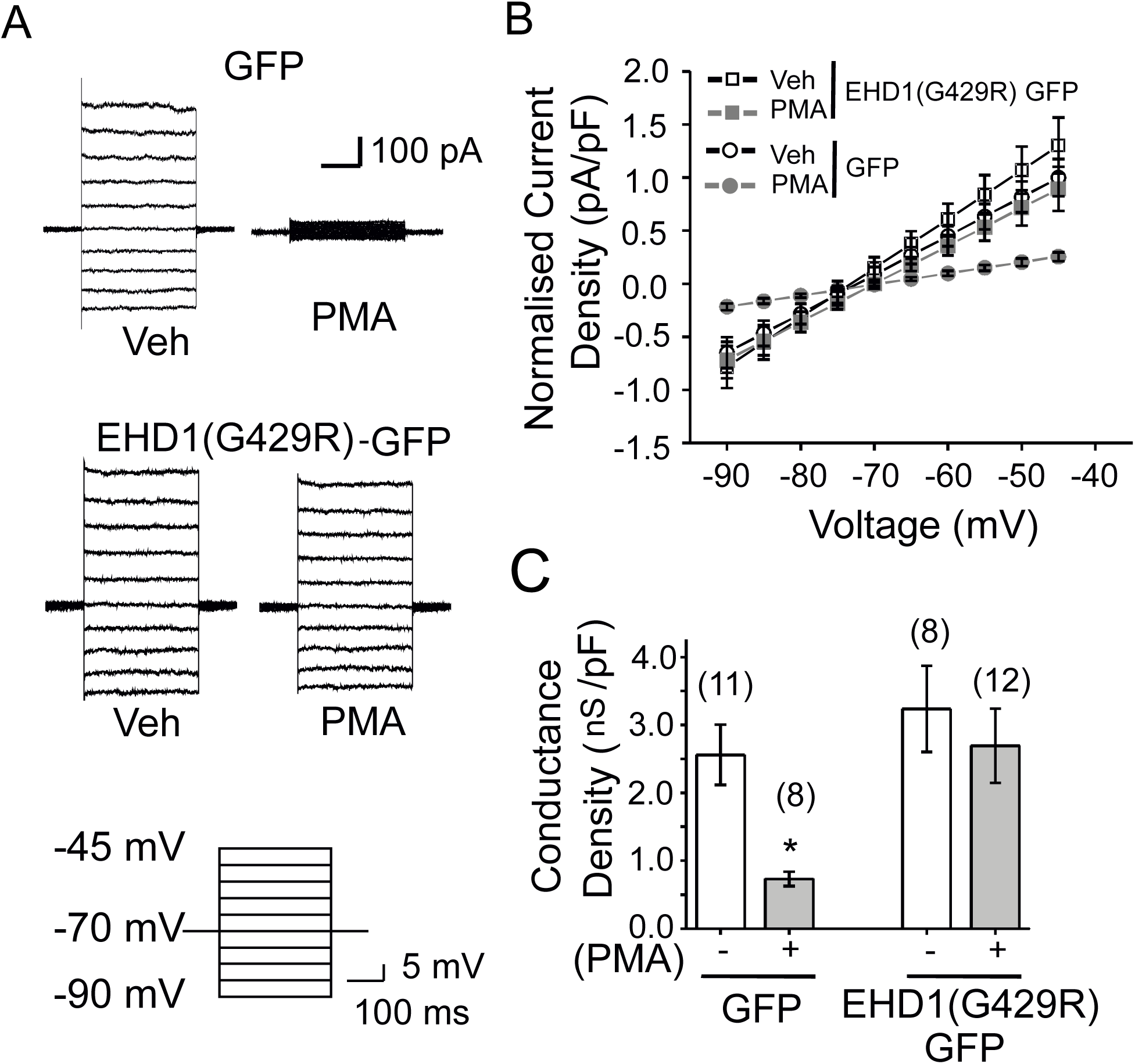
EHD-1(G429R)-GFP prevents PKC-induced downregulation of K_ATP_ channel current density in the INS-1 832/13 pancreatic β-cell line. Cells expressing either GFP or EHD-1(G429R)-GFP were treated for 30 min with vehicle (Veh, 0.1 % DMSO) or 100 nM PMA and subjected to whole-cell voltage-clamp recording. (A) Representative current traces recorded during 5 mV pulses from −90 mV to −45 mV from a holding potential of −70 mV (protocol at the bottom). (B) Mean ± SEM of current density-voltage relationships from recordings performed as in (A); the data were normalised to the current density at −45 mV of GFP transfected cells. (C) Mean ± SEM of conductance density measured from the slopes of curves in (B); * indicates significance (*p* < 0.05).

### PKCε is responsible for downregulation of K_ATP_ channels

PKC comprises a family of 12 closely related enzymes, classified into 3 groups: classical (α,β,γ), novel (δ,ε,η,θ) and atypical (ζ, ι) (Steinberg, 2008)(Wu-Zhang and Newton, 2013). Many of these isoforms are expressed in β-cells (Warwar et al., 2006). We sought to identify the PKC isoform responsible for PMA-induced downregulation of K_ATP_ channels using pharmacological inhibitors and dominant negative constructs of PKC. A role for atypical PKC isoforms was ruled out because they are PMA insensitive (Wu-Zhang and Newton, 2013). We excluded a role for classical PKC, because PMA-induced endosomal accumulation of the channel was abolished by the broad spectrum PKC inhibitor, chelerythrine, but not by GÖ6976, an inhibitor selective for classical PKC (Wu-Zhang and Newton, 2013) (Figure 5A). These data narrow down the PKC responsible for K_ATP_ channel downregulation to a novel isoform.

**Figure 5.**
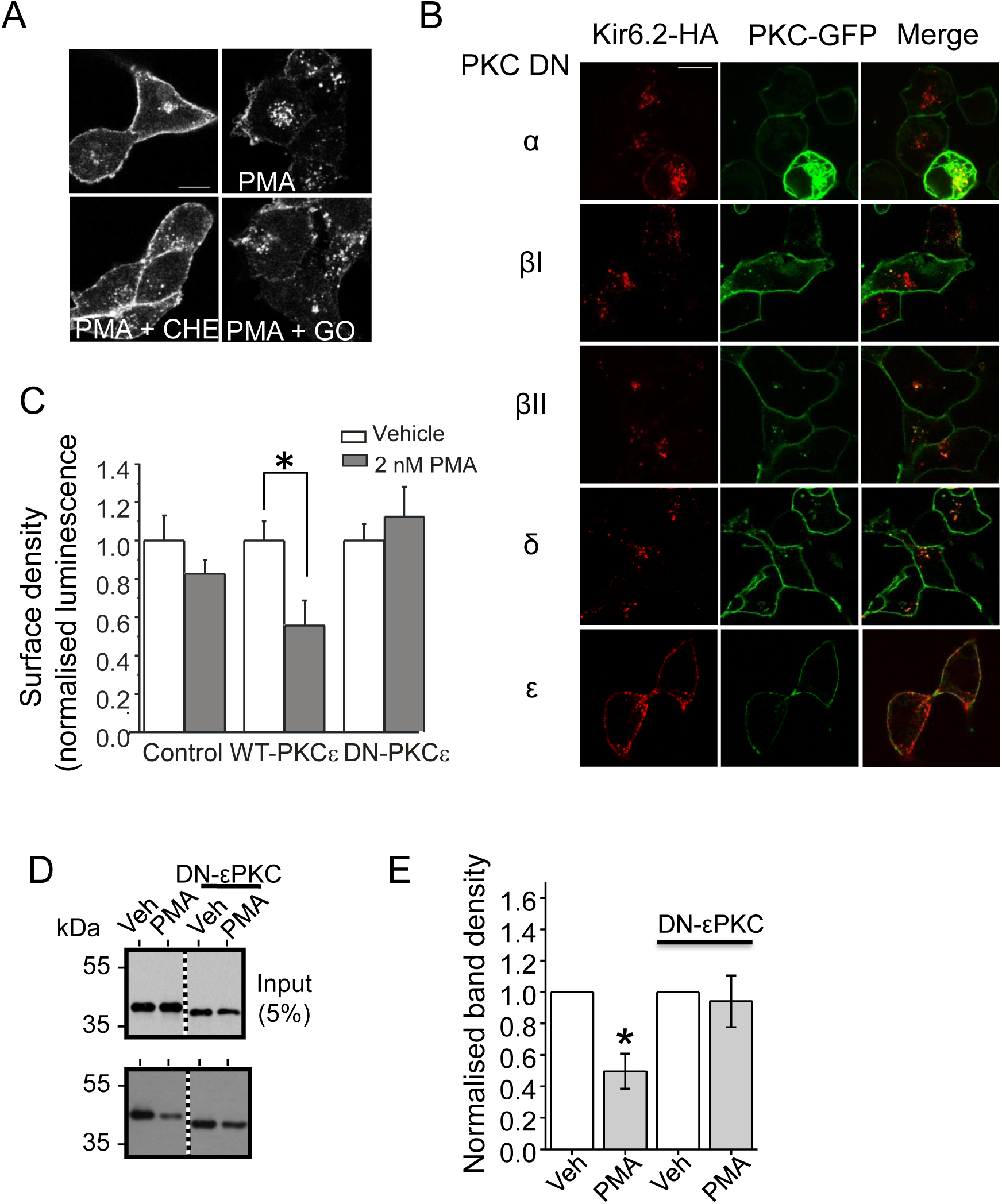
PKCε promotes degradation and downregulation of K_ATP_ channels. (A) Chelerythrine, but not GÖ6976, prevents PMA-induced cellular distribution of surface labelled K_ATP_ channels. HEK 293 cells expressing the recombinant channels were pre-treated with chelerythrine (10 µM) or GÖ6976 (1 µM) or vehicle (0.1 % DMSO) and incubated for 1 h with anti-HA antibodies in the presence of PMA (100 nM) and the continued presence of inhibitors. Representative images (from 3 independent experiments) of cells stained with secondary antibodies are shown; scale bar 10 µm. (B) Dominant negative PKCε blocks PMA induced K_ATP_ channel redistribution. HEK 293 cells expressing the recombinant channels were transfected with the indicated GFP-tagged dominant negative (DN) constructs of PKC isoforms. Cells were treated with PMA (100 nM) and anti-HA antibodies for 1 h prior to permeabilisation and staining with Cy3 conjugated secondary antibodies. Representative images (from 3 independent experiments) are shown; scale bar 10 µm. (C). Expression of PKCε downregulates K_ATP_ channels. HEK 293 cells stably expressing the recombinant channels were either not transfected (control) or transfected with the indicated PKCε constructs. Cells were treated with the vehicle (0.1% DMSO) or PMA (2 nM, 1 h) and the surface density measured as described in methods. Mean ± SEM (n=3) of surface density normalised to respective controls are shown. (D-E) Overexpression of DN-PKCε prevents PMA-induced degradation of K_ATP_ channels. Experiment was performed as described in the legend to Figure 2C. Representative blots (D) and the corresponding mean ± SEM data (n=3) (E) are shown. * Indicates significant difference (*p* < 0.05).

To underpin the PKC isoform responsible for the PMA effects, we transfected the HEK-293 cells stably expressing HA-tagged K_ATP_ channels with dominant negative constructs of GFP-tagged novel PKC isoforms (δ and ε). We have also used dominant negative classical PKC (α, βI and βII) isoforms as negative controls. PMA treatment led to the activation of all transfected isoforms as evident from their distribution to the plasma membrane (green fluorescence, Figure 5B). However, with the exception of the dominant negative PKCε, none of the other constructs prevented the ability of PMA to induce endosomal accumulation of the channel. These data support a role for PKCε in the endosomal retention of K_ATP_ channels. To demonstrate that endosomal retention is the reason for channel downregulation, we transfected the cells with the wild-type and dominant negative PKCε and determined the PMA effect on the channel surface density. In this experiment, we have used a low concentration (2 nM) of PMA to minimise the activation of endogenous PKC. The results (Figure 5C) show that at this dose, PMA had no effect on non-transfected cells, but significantly reduced the surface density of channels in cells transfected with the WT-PKCε (*p*<0.05), but not the dominant negative PKCε (*p*>0.05).

We next wanted to confirm that PMA-induced channel degradation (Figure 2) is due to activation of PKCε. Using the biotinylation-based assay, we show that the dominant negative PKCε was able to fully rescue PMA-induced degradation of endocytosed K_ATP_ channels (Figure 5D-E). Taken together, we conclude that activation of PKCε causes downregulation of plasma membrane K_ATP_ channels by promoting their endosomal degradation.

## Discussion

A recent study demonstrated that HG triggers β-cell excitation by downregulating the cell surface K_ATP_ channels (Han et al., 2018). The authors reported that the HG-induced downregulation occurs by PKC-induced endocytosis. However, other studies showed that PKC does not affect endocytosis, but prevents recycling, while promoting channel degradation (Mankouri et al., 2006; Manna et al., 2010). To address this discrepancy, here we have determined the mechanistic details of channel downregulation using PKC-resistant mutant channels. We demonstrate that (i) mutation of the dileucine trafficking motif (^355^LL^356^) located on the Kir6.2 subunit of the channel prevents PKC-induced endosomal degradation and downregulation, (ii) PKC-induced channel degradation and downregulation are mediated by EHD proteins, and (iii) the PKC isoform responsible for the channel degradation is PKCε. We conclude that endosomal degradation, but not endocytosis, underlies PKC-induced downregulation of cell surface K_ATP_ channels. We discuss the implications of these findings for GSIS.

### L355P mutation in Kir6.2 abolishes the ability of PMA to alter post-endocytic trafficking of K_ATP_ channels

Dileucine motifs play diverse roles in membrane protein trafficking including in endocytosis, recycling and lysosomal trafficking (Bonifacino and Traub, 2003). Kir6.2 has a dileucine motif (^355^LL^366^) that has been shown to play a role in PKC-mediated channel downregulation (Hu et al., 2003; Mankouri et al., 2006). Although an earlier study reported that the dileucine motif in Kir6.2 promotes channel endocytosis, mutation of the motif to AA (LL/AA) (Mankouri et al., 2006), Figure 1G) or proline (L355P) (Figure 1 C-D) failed to prevent endocytosis. Instead, these mutations abolished the ability of the PKC activator PMA to inhibit recycling (Figure 1E-F, H). This was accompanied by a loss in the ability of PMA to induce lysosomal trafficking (Figure 2A-B), degradation (Figure 2C), and downregulation (Figure 2D) of plasma membrane K_ATP_ channels. These results demonstrate that the dileucine motif determines PKC regulation of cell surface density of K_ATP_ channels by affecting channel recycling and /or degradation. Knock-in studies are required to determine if the functionally silent L355P human mutation (Sakura et al., 1996) is associated with type 2 diabetes by virtue of its ability to prevent PKC-dependent K_ATP_ channel downregulation. Regardless of the pathophysiological relevance, from the cell biological perspective, our results raise the key question of whether reduced recycling leads to increased channel degradation or vice versa.

### EHD proteins mediate PKC-induced degradation and downregulation of K_ATP_ channels

The above question promoted us to first examine the mechanism by which PKC regulates K_ATP_ channel recycling. EHD-1 is known to play a major role in recycling of many membrane proteins (Naslavsky and Caplan, 2011; Park et al., 2004; Picciano et al., 2003). Consistent with a role for EHD proteins, co-immunostaining experiments revealed PMA to increase the localisation of internalised channels to EHD-positive endosomes (Figure 3A). However, the dominant negative EHD-1(G428R)-GFP, which is known for its ability to inhibit recycling of membrane proteins (Naslavsky and Caplan, 2011; Park et al., 2004; Picciano et al., 2003), failed to inhibit K_ATP_ channel recycling. On the contrary, it rescued PMA-induced inhibition of channel recycling (Figure 3B-C). To explain this unexpected finding, we hypothesised that the dominant negative EHD-1 prevents PMA-induced degradation of the channel and thereby sustain the endosomal pool of channels available for recycling. Consistent with this idea, EHD-1(G428R)-GFP prevented PMA-induced channel degradation (Figure 3D), and led to increased recycling (Figure 3C). These findings ruled out a direct role for EHD-1 in PKC regulation of recycling; instead they support the idea that EHD-1 indirectly reduces recycling by diverting the pool of recycling endosomes to degradation, and thereby downregulates the number of channels at the plasma membrane. Consistent with the cell biological and biochemical data (Figure 3), whole-cell patch clamp experiments on the INS-1 cell line demonstrated that EHD-1(G428R)-GFP abolished the ability of PMA to downregulate K_ATP_ channel conductance in INS1 832/13 cells (Figure 4).

While this finding is of considerable interest, we do not know how EHD-1, known for its role in recycling, plays a role in PKC-induced channel degradation. Recent studies suggest that EHD-1 can form heterodimers with other members of the family capable of mediating trafficking to late endosomes/lysosomes (Naslavsky and Caplan, 2011). Intriguingly, the effect of EHD-1(G428R)-GFP is strikingly similar to that of L355P mutation: both caused an increase in channel degradation, reduced recycling and promoted downregulation (Figures 1-3). Thus there appears to be a mechanistic link between the dileucine motif and EHD proteins in mediating PKC-induced degradation of K_ATP_ channels, which could be important for β-cell excitation. Further studies are required to elucidate this intriguing relationship.

### Cell biological mechanism underlying PKC-induced K_ATP_ channel downregulation

Han et al have recently reported that HG downregulates cell surface K_ATP_ channels in INS-1 cells, implicating a major role for this mechanism in insulin secretion (Han et al., 2018). They suggested that the HG-induced downregulation is due to increased endocytosis triggered by PKC activation. The basis for this conclusion was the finding that dynasore, an inhibitor of endocytosis, prevents HG-induced channel downregulation. Here we explain how the dynasore effect could be misinterpreted in the absence of careful consideration of other aspects of endosomal traffic. Acute regulation of cell surface density of K_ATP_ channels (Manna et al., 2010), like most membrane proteins (Bonifacino and Traub, 2003), is determined not just by endocytosis, but by a balance between endocytosis, recycling and lysosomal degradation. In the control conditions, K_ATP_ channels undergo rapid cycles of endocytosis and recycling, but their rate of degradation is very slow (Manna et al., 2010). In the PKC-stimulated conditions, endocytosis and recycling continues to occur normally, but the rate of channel degradation increases markedly (Manna et al., 2010). This is expected to lead to a decline in the endosomal pool of K_ATP_ channels available for recycling, and thus to their downregulation at the plasma membrane. In this scenario, in the presence of dynasore, any channel that recycles back is expected to be retained at the plasma membrane. Since recycling occurs much faster than lysosomal trafficking (Manna et al., 2010), dynasore treatment is expected to cause a rapid decline in the endosomal pool of channels and a concomitant increase at the cell surface. Such an effect will appear as though dynasore had inhibited PKC-induced endocytosis. We therefore propose a reinterpretation of Han et al’s observation: HG causes downregulation of K_ATP_ channels by increasing their endosomal degradation (coupled with reduced recycling), rather than promoting endocytosis.

### PKCε is responsible for endosomal degradation and downregulation of K_ATP_ channels

Pancreatic β-cells express several isoforms of PKC, which are thought to play different roles in GSIS (Trexler and Taraska, 2017). They are activated during high glucose conditions and influence insulin exocytosis by activating components of the exocytotic machinery. However, much less is known about the PKC isoforms involved in trafficking of ion channels regulating GSIS. We therefore sought to identify the PKC isoform responsible for K_ATP_ channel downregulation. Using pharmacological agents (Figure 5A) and dominant negative (kinase-dead) constructs (Figure 5B), we demonstrated that the isoform responsible for K_ATP_ channel downregulation (Figure 5C) and degradation (Figure 5D-E) as PKCε.

### Possible patho-physiological significance of the current findings

We consider the potential implications of PKC regulated K_ATP_ channel trafficking for GSIS. GSIS occurs in two phases: a fast, transient first phase (∼1-10 min) followed by a slow, sustained second phase (∼10 to 60 min). The first phase is dependent on the closure of K_ATP_ channels, whereas the second phase is thought to be K_ATP_ channel-independent. This notion is supported by the evidence that activation of K_ATP_ channels with diazoxide (Straub and Sharp, 2002), or V59M genetic mutation (Girard et al., 2009), abolishes the first, but not the second phase of GSIS.

The second phase is associated with an increased production of DAG (Wolf et al., 1990) and activation of several PKC isoforms (α,β,ε and θ) in the β-cell (Trexler and Taraska, 2017; Warwar et al., 2006). The different PKC isoforms appear to target different components of machinery associated with insulin secretion (Trexler and Taraska, 2017). In relation to the K_ATP_ channel, an yet-to-be identified PKC isoform activates the channel via threonine180 phosphorylation at Kir6.2 (Light et al., 2000). Although not demonstrated in β-cells, this could compromise the biophysical ability of ATP to close the channels during the second phase of GSIS. In order to sustain depolarisation during the second phase, it appears that β-cells have adopted a cell biological mechanism, whereby K_ATP_ channels are removed from the cell membrane (Han et al., 2018) and subjected to lysosomal degradation (Manna et al., 2010). This study shows that the latter effect is mediated by PKCε (Figure 5). Although we currently do not have direct evidence, there are several published studies that could support the idea that PKCε-mediated downregulation of K_ATP_ channels may underlie the second phase of GSIS. First, HG causes β-cell excitation by triggering PKC-dependent downregulation of K_ATP_ channels (Han et al., 2018). Second, knock-down of PKCε abolishes the second, but not the first, phase of GSIS (Warwar et al., 2008). Thirdly, islets of Goto-Kakizaki (GK) diabetic rats have reduced levels of PKCε compared with normal mice (Warwar et al., 2006). Finally, GK diabetic rats display a marked loss of the second, but not the first, phase of insulin secretion (Portha et al., 1991). Once the secreted insulin restores the glucose levels back to normal, [AMP]+[ADP]/[ATP] ratio in the cell increases, leading to AMPK (AMP-dependent protein kinase) activation (Lim et al., 2009; Salt et al., 1998). AMPK activation stimulates expression of K_ATP_ channels (Lim et al., 2009; Smith et al., 2006), restoring the channel density back to normal to prevent unwarranted insulin secretion.

In summary, our model suggests that HG stimulates the first phase of insulin secretion through ATP inhibition of channel opening (Po), and the second phase via PKCε-induced downregulation of the number (N) of K_ATP_ channels at the cell surface. Thus we propose that K_ATP_ channels regulate not only the first, but also the second phase of GSIS through fundamentally distinct mechanisms (Figure 6). Since the amount of insulin secreted during the second phase is far greater than that of the first phase, future anti-diabetic therapeutics should aim to target K_ATP_ channel downregulation.

**Figure 6.**
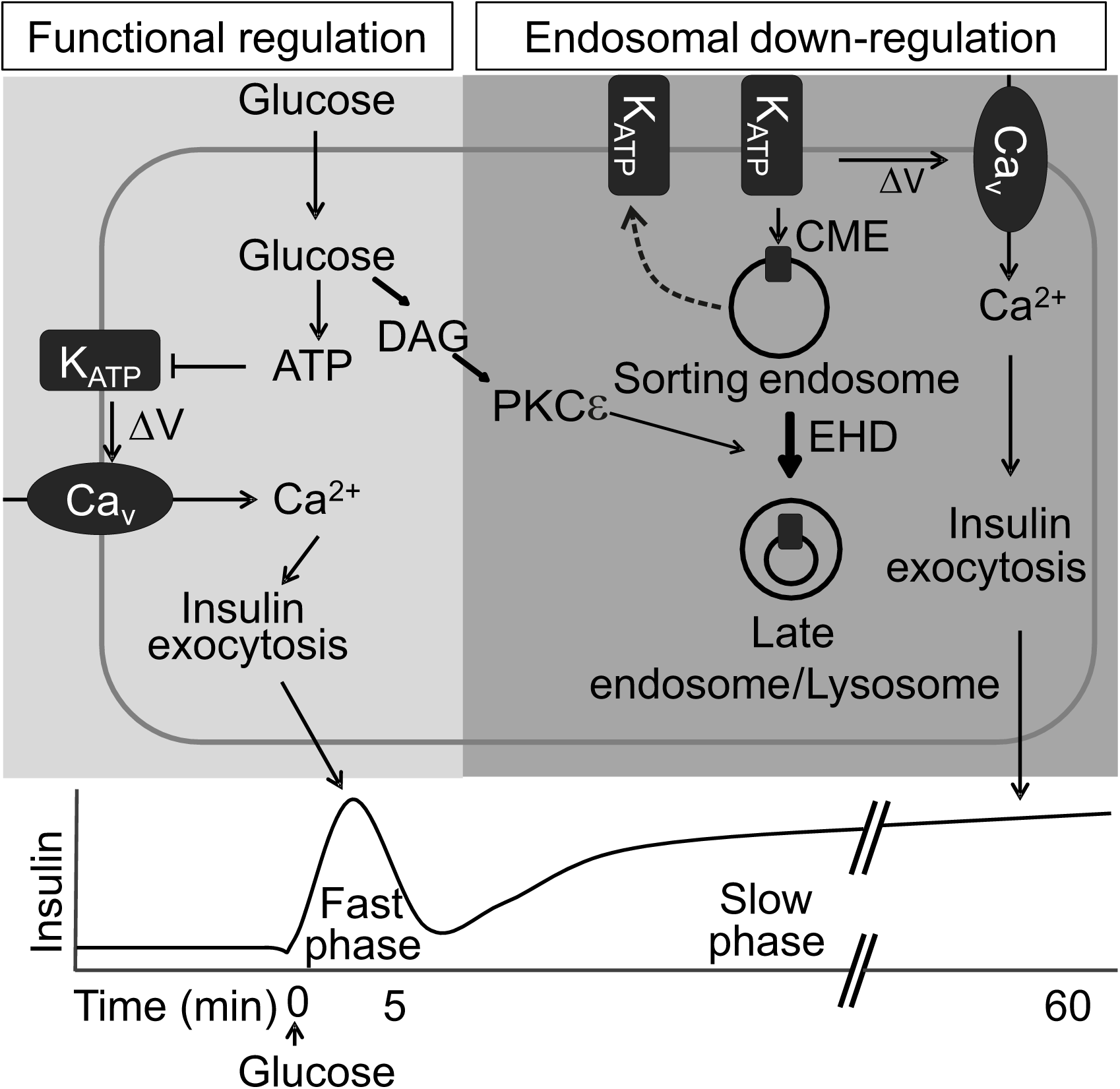
Schematic of the proposed mechanisms by which K_ATP_ channels regulate the two phases of GSIS. Glucose metabolism generates ATP and DAG. ATP inhibits the K_ATP_ channels (decreases Po), leading to Ca^2+^ entry and insulin exocytosis that comprises the fast phase of GSIS. This functional regulation is shown on the left (shaded in light grey). DAG activates PKCε leading to the trafficking of constitutively endocytosed channels to late endosomes/lysosomes (bold arrow) where they are degraded. The PKC-induced degradation is mediated by EHD proteins and requires the dileucine motif of Kir.6.2. Degradation reduces the endosomal pool of channels, and thereby indirectly reduces recycling (indicated by broken line arrow). This results in the downregulation of the number of K_ATP_ channels (N) at the plasma membrane, contributing to β-cell depolarisation, Ca^2+^ entry and insulin exocytosis during the second phase of GSIS. The DAG effect on endosomal downregulation of the channel is shown on the right (dark grey).

## Acknowledgements

This work was supported by the Medical Research Council, UK ((G0802050) and the Overseas Research Students Award Scheme (RK). We thank Dr JD Lippiat for helpful suggestions.

## Materials and methods

### Molecular biology and Cell culture

pCDNA3-Kir6.2-HA and pCDNA6-SUR1 constructs are as described previously (Manna et al., 2010). Kir6.2 contains the HA (haemaglutinin A) epitope in the extracellular portion of the protein. INS-1 832/13 cells (kindly provided by C.B. Newgard, Duke University School of Medicine) and HEK-293 cells stably expressing the K_ATP_ channels were cultured as described (Manna et al., 2010). EHD-1(G429R)-GFP was a gift from F. R. Maxfield (Weill Cornell Medical Collage). pXLIN-PKCα-EGFP-K368R and pEGFP-N1-PKCβI-K371R were provided by Dr. M. Breune (Dresden), pBK-CMV-PKCβ2-K371R-GFP by Dr. Y. Hannun (Medical University of South Carolina), pEGFP-N1-PKCδ-K371R by Dr. M. Reyland (University of Colorado Denver), and pEGFP-N1-PKCε-K437R by Dr. J. W. Soh (Inha University, Korea). Mutations were introduced by the QuikChange method. Cells were grown on coverslips or in dishes and transfected using FuGENE® 6 (Promega).

### Immunofluorescence

To detect HA-tagged channels at the cell surface, cells were fixed in 2% paraformaldehyde (PFA) for 15 min, washed with PBS (phosphate buffered saline) and blocked with 1% ovalbumin, 10 mM HEPES, DMEM, pH 7.4 for 30 min. Cells were then incubated with rat anti-HA antibodies (Roche, 3F10; 0.2 μg/ml) for 1 hr, washed with PBS (3 times) and stained with Cy3-conjugated donkey anti-rat IgG (Jackson ImmunoResearch Laboratories; 1.25 µg/ml). To detect total channels, PFA-fixed cells were permeabilised with a 1:1 mixture of methanol and acetone (−20 °C) and then stained.

To detect Internalised channels, cells were incubated with rat anti-HA antibodies for 30 min at 4° C, washed with PBS (3 times), and then incubated at 37 °C to permit internalisation of antibody labelled channels. In some experiments, cells were incubated with rat anti -HA antibodies at 37 °C for 30 min to allow continuous internalisation. Internalised channels were stained after fixing and permeabilisation as outlined above.

Recycling of internalised channels back to the cell surface was detected as described before (Manna et al., 2010), with minor modifications. Briefly, cells were incubated with rat anti-HA antibodies at 37 °C for 1 hr to allow internalisation. After three PBS washes, cells were incubated at 14 °C (a temperature not permissible for recycling) with unconjugated goat anti-rat IgG (Sigma, 4 μg/ml) to block labelled channels remaining at the cell surface. After washing with PBS (3 times), cells were incubated with Cy3-conjugated donkey anti-rat secondary antibodies (1.25 µg/ml) at 37 °C for 30 min to stain anti-HA labelled channels returning to the cell surface.

Intracellular compartments were labelled with either mouse anti-CD63 (abcam, NK1/C3, 1-10 μg/ml) or mouse anti-Rme-1-4 (Santa Cruz Biotechnology, EHD (E-5), 0.8 μg/ml) and stained with Cy5-conjugated donkey anti-mouse IgG (Jackson ImmunoResearch Laboratories; 1.25 µg/ml).

Cells were imaged using an upright Axiovert LSM 510 confocal microscope (Zeiss) fitted with *a* 63x-1.4NA oil DIC objective using a helium/neon laser (Ex. 550 nm, Em. 570 nm for CY3; Ex. 625 nm, Em. 670 nm for CY5) or an argon laser (Ex. 488 nm, Em. 512 nm for GFP). Representative images from three or more independent experiments are shown with 10 µm scale bars. Colocalisation was analysed using Image J (NIH) and expressed as Pearson correlation coefficient.

### Chemiluminescence

Quantification of channel density, rates of endocytosis and recycling were performed by the chemiluminescence assay as described previously (Manna et al., 2010). The ratio of surface to total channel density and channel internalisation were measured as described by Taneja et al (Taneja et al., 2009) and Manna et al (Manna et al., 2010) respectively. The labelling steps are similar to those described above, except that HRP-conjugated goat anti-rat IgG (Sigma, 1-8 µg/ml) was used in place of fluorescent secondary antibodies. The labelled cells were lysed and HRP binding to channels was quantified by measuring the chemiluminescence emitted from the HRP-catalysed oxidation of Lumigen® PS*-*atto (Lumigen, Inc) ECL substrate. The chemiluminescence was normalised to the protein concentration of the lysates, measured by the bicinchoninic acid assay.

To quantify recycling, cells were incubated with rat anti-HA primary antibodies at 37 °C for 1 hr to allow internalisation. After three PBS washes to remove primary antibodies, cells were incubated with goat HRP-conjugated anti-rat IgG at 37 °C for 0 or 30 min (plus or minus 100 nM PMA/0.1% DMSO) to label channels recycling to the cell surface. The cells were washed with chilled PBS three times prior to measurement of HRP activity as described above. The difference between the 30 and 0 min values was taken as a measure recycling.

### Degradation of biotinylated surface channels

For this, a biotinylation assay was used (Manna et al., 2010). Surface proteins were labelled with EZ-Link Sulfo*-*NHS-LC-Biotin (Thermo Scientific, 0.3 mg/ml in PBS) at 4 °C for 30 min. After quenching excess reagent with ice-cold 50 mM glycine/PBS, cells were incubated at 37 °C for 2 hrs in serum free DMEM in the absence or presence of PMA, before lysing in lysis buffer ((1% Triton X100, 50 mM HEPES, 150 mM KCl, 1 mM MgCl_2_, pH 7.4, with 1x EDTA-free protease inhibitors, Roche) at 4 °C for 2 hrs. The lysates were centrifuged (15,000 x*g*, 30 min 4 °C) and the supernatants mixed with 30 μl of NeutrAvidin agarose beads (Thermo Scientific) for 2 hrs at 4 °C to adsorb the biotinylated proteins. The beads were washed with lysis buffer and subjected to Western blotting. HA-Kir6.2 was detected with HRP-conjugated rat anti-HA antibody (Roche, 3F10, 0.005 μg/ml). GFP was detected with mouse anti-GFP antibodies (AbCam; 0.2 μg/ml) and HRP-conjugated anti-mouse IgG (Sigma; 1.0 μg/ml). 5% of lysates (input) were analysed in a parallel lane on the same blot. Band densities were measured using Image J (NIH); intensities of biotinylated Kir6.2 bands (surface protein) were normalised to input band (total cellular Kir6.2) densities.

### Electrophysiology

Currents through K_ATP_ channels were recorded in the standard whole cell configuration as described previously (Manna et al., 2010). INS-832/13 cells grown on coverslips were transfected with GFP or EHD-1(G429R)-GFP. Prior to recording, cells were incubated in serum-free RPMI for 30 min at 37 °C, followed by further 30 min-incubation in the presence of vehicle (0.1% DMSO) or 100 nM PMA. Patch pipette were made from thin walled glass pipettes (∼2.5 MΩ) that were filled with intracellular solution (100 mM KCl, 40 mM KOH, 1.7 mM MgCl_2_, 1 mM CaCl_2_, 10 mM EGTA, 10 mM HEPES, pH 7.2). Currents were recorded in extracellular solution (138 mM NaCl, 5.6 mM KOH, 2.6 mM CaCl_2_, 1 mM MgCl_2_, 10 mM HEPES, pH 7.4), using an EPC10 amplifier with PatchMaster (HEKA). Current densities (pA/pF) were normalised to the GFP expressing, vehicle (control)-treated recordings and plotted against voltage (mV). Conductance density (nS/pF) was calculated for each recording from the slope of the current-voltage relationship.

### Data analysis

Data were analysed using Origin 7. Results are expressed as mean ± SEM of ‘n’ (shown in figures) independent experiments; * indicates significance at *p* < 0.05, determined using the two-tailed Student’s t-test.

